# Exposure to naturalistic occlusion promotes generalized, human-like robustness in deep neural networks

**DOI:** 10.64898/2026.04.23.720370

**Authors:** David D Coggan, Frank Tong

## Abstract

Human object recognition is robust to challenging conditions, such as when one’s view of an object is fragmented due to an occluding foreground object. In comparison, deep neural networks (DNNs) are typically more susceptible to occlusion, suggesting that human vision relies on distinct mechanisms. Here, we investigated the role of visual diet in the emergence of these mechanisms by asking whether human-like robustness might arise in DNNs when trained with image datasets that better reflect the properties of occlusion in natural vision. We trained convolutional and transformer DNNs to classify clear images only, images augmented with artificial occluders (i.e., geometric shapes) or natural occluders (objects segmented from photographs). We then evaluated DNN occlusion robustness and compared their performance profiles with 30 human participants. We found that DNNs trained with artificial occluders remained vulnerable to natural occlusion and exhibited less human-like performance than those trained with natural occlusion. Our findings suggest that human robustness to visual occlusion arises from learning to disentangle natural objects from each other rather than simply learning to recognize objects from partial views. They also imply that commonly used forms of artificial occlusion are unsuitable for the evaluation or promotion of robustness to real-world occlusion in DNNs.

## INTRODUCTION

For over a decade, human object perception has been best approximated by a class of image-computable vision models known as deep neural networks (DNNs). Many models within this class can perform visual tasks such as object classification with roughly human-level competence ^1^ while strongly predicting human behavioral responses and underlying neural representations ^2–15^. However, compared to humans, DNN performance is generally much poorer if images are naturally low-quality ^16^ or have been deliberately perturbed ^17–24^, suggesting that human object perception involves different or additional mechanisms not currently instantiated in DNNs.

Previous studies have shown that DNNs become more robust and human-aligned when training images are perturbed by applying noise ^20,25^ or blur ^21,26,27^, although generalization to untrained perturbations might be limited ^19,28^. Insofar as these augmentations correct for under-represented aspects of human visual experience in standard training datasets, such findings suggest that humans acquire robust vision by learning from their extensive exposure to challenging viewing conditions throughout development – a concept that we term the *ecological vision hypothesis*. However, the extent to which human robustness to occlusion might depend on visual experience has not been explored, as previous computational studies have largely focused on architectural modifications for improving occlusion robustness in DNNs ^17,22,24,29–34^.

However, the goal of instantiating human-like robustness to occlusion in DNN models by modifying their training data might not be as straightforwardly achievable because, unlike blur or noise, occlusion has unique characteristics of introducing complex signals from the foreground distractor object, which encompasses a vast, high-dimensional space of possible perturbations. One might hypothesize that DNNs exposed to any form of additional occlusion augmentation will exhibit more human-like behavior, in which case human robustness could be attributed to the benefits of extensive training with partial views of objects, irrespective of the nature of the occluder itself. Alternatively, human-like robustness might only emerge when the training occluders are naturalistic, i.e., constituted by real objects, which would imply that human robustness is specialized for disentangling natural objects that frequently obscure one another. A final possibility is that the training of DNNs with augmented examples of partially occluded objects might prove inadequate for attaining human-like recognition of occluded objects, which would imply that human robustness to occlusion might not be readily explained by visual diet alone.

To investigate how generalized, human-like robustness to occlusion might be induced in DNNs, we developed the Visual Occluders Dataset (https://github.com/ddcoggan/VisualOccludersDataset), a novel dataset of occluder masks that can be applied to object images during training and evaluation. Using ImageNet-1K, DNNs were trained with either clear images only or with images augmented with one of two sets of computer-generated, silhouette occluders (*artificial 1*, *artificial 2*); natural occluders obtained by segmenting real objects from photographs (*natural*); or the natural occluders rendered as silhouettes (*natural silhouette*). Examples of the occluder training sets can be seen in **Figure 1a**. After training, we evaluated DNN robustness on a separate image validation set while applying each occluder set used in training. To evaluate DNN-to-human alignment, we conducted a separate human behavioral experiment (N=30) by applying a subset of occluders (those in *artificial 1* and *natural silhouette*) to eight classes of objects (shown in grayscale to remove obvious color cues), and then compared classification accuracies (8-AFC) between DNN models and human observers.

**Figure 1.**
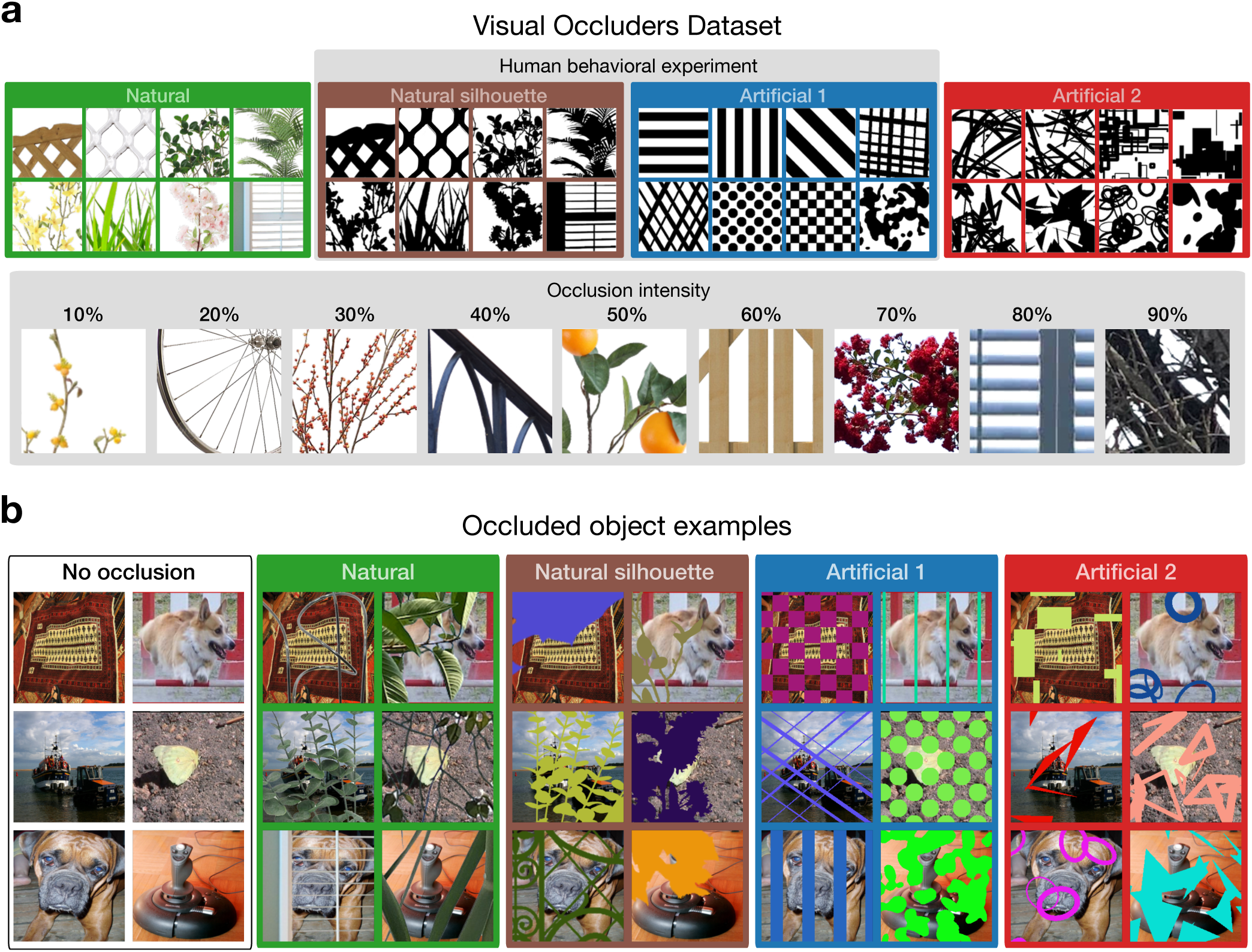
The Visual Occluders Dataset. (**a**) Examples of occluders used in the 4 DNN occlusion training conditions (top row). Bottom row shows natural occluders that vary in occlusion intensity based on the proportion of the target image that will be obscured. The dataset contains up to 1000 individual occluder masks for each occluder type and intensity. For the purposes of the present study, 16 of the 30 artificial occluder types contained in the dataset were arbitrarily selected and allocated to artificial sets 1 or 2. (**b**) Examples of occluded object images used for each DNN training condition. For clarity, images are presented without subsequent preprocessing transformations that were used during training (e.g., color jitter, weak randomized levels of Gaussian blur, normalization, etc., see **Methods**).

We found that, compared to baseline models trained without occlusion, DNNs trained with artificial forms of occlusion performed poorly at generalizing to objects with natural occluders and exhibited a divergent profile of performance accuracy across occluder types when compared with human observers. By contrast, naturalistic occlusion training led to more generalized robustness across a wide range of occlusion conditions that strongly predicted human performance accuracy. These findings provide support for our ecological vision hypothesis, namely that humans acquire robustness to occlusion through prevalent encounters with a variety of diverse foreground objects in everyday vision, allowing the visual system to learn how to discount these complex distracting signals.

## RESULTS

### Occlusion robustness

A separate set of DNN models were trained on each of five possible visual diets: no occlusion, natural, natural silhouette, artificial 1 or artificial 2 (see **Figure 1**). For the occlusion-trained models, the amount of occlusion applied to individual images from the ImageNet-1K training set varied from 0% to 50% occlusion. After training, each DNN model was evaluated with all five occlusion conditions using the ImageNet-1K validation set. For each DNN, we first measured top-1 classification accuracies for each occluder set and intensity in the Visual Occluders Dataset. We then aggregated these scores across DNN architectures and occlusion intensities to obtain a single accuracy score for each pairwise train / test combination across the five occlusion conditions. As can be seen in **Figure 2a**, all DNN models performed best in the no occlusion condition, and when faced with occluded object images, the occlusion-trained models performed best on the test occluders that matched those used during training (i.e., diagonal cells indicated in green). Outside of the diagonal, however, it is evident that DNNs trained with either artificial occlusion or natural silhouette performed poorly at generalizing to images with natural occlusion. By contrast, DNN models trained with natural occlusion showed comparatively better generalization to both artificial and natural silhouette occluders.

**Figure 2.**
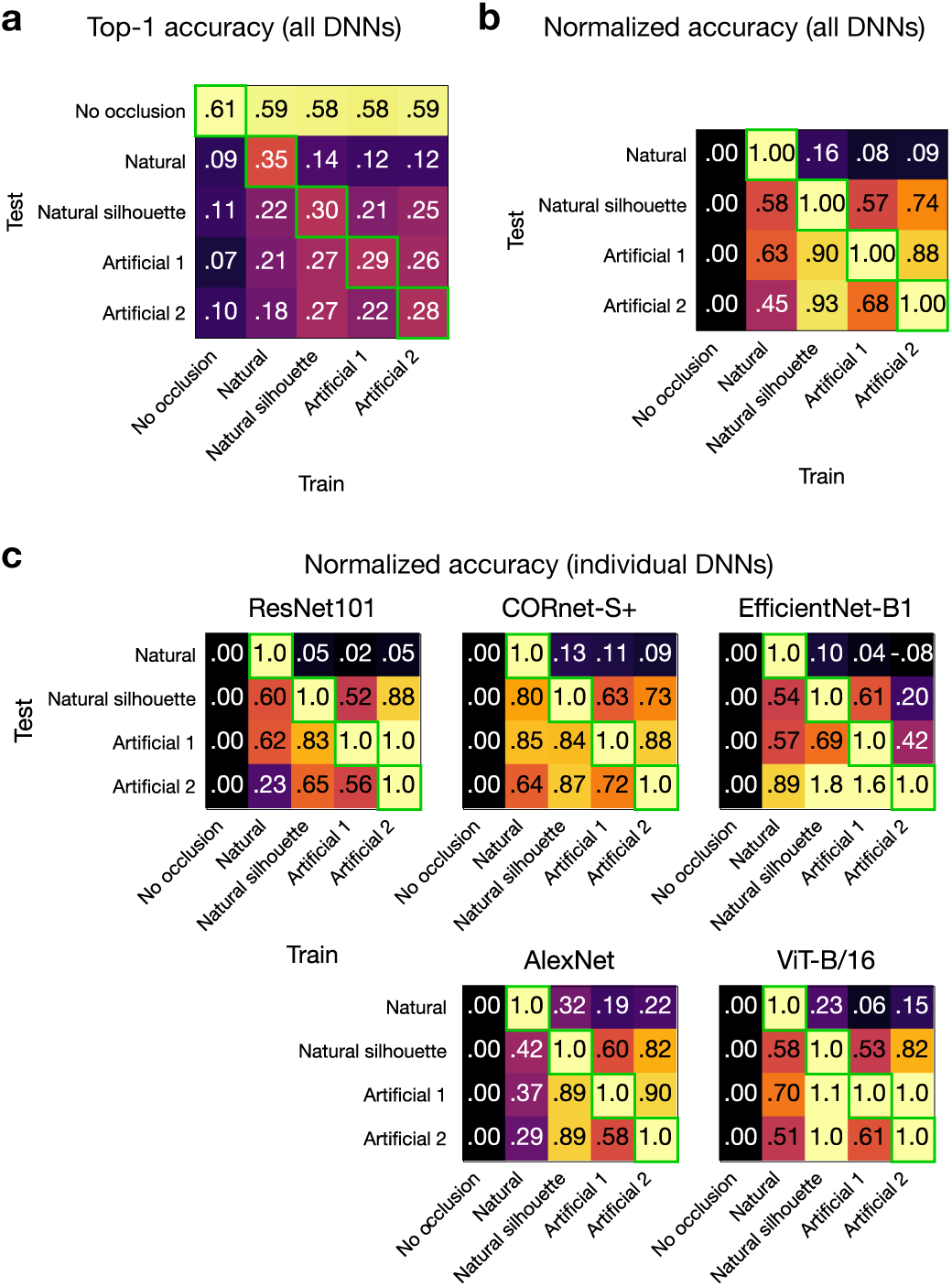
Occlusion robustness in DNNs trained with different forms of occlusion, measured using top-1 classification accuracy for object images drawn from the ImageNet-1K validation set. For each occlusion intensity (10-90%), DNNs were tested on the entire validation set applied with a given occluder condition, with ‘silhouette’ style occluders imposed using a randomly chosen color in each instance (as in Figure 1b). (**a**) Mean classification accuracy plotted by DNN training condition and occluder test condition, averaged across all occlusion intensities and DNN architectures. Green boxes indicate when DNNs were trained and tested on the same condition. (**b**) Accuracy scores from (a) were normalized based on the range of performance expected for each set of test occluders (see **equation 1**). (**c**) Normalized accuracy scores, as in (b), but shown separately for each individual DNN architecture.

Because overall accuracy varied somewhat across the different test occluder conditions, we further inspected the results after normalizing the accuracy scores. Specifically, we calculated normalized accuracy scores (*S*) by rescaling raw accuracies (*A*) based on floor levels of performance in DNNs trained without occlusion (*B*) and ceiling levels in DNNs trained on the corresponding test occluder condition (*C*), using the following equation:

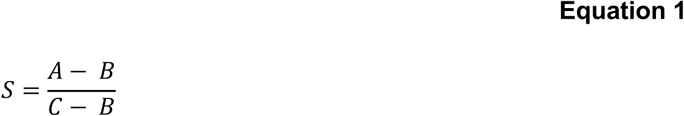

Thus, a score of zero indicates that DNNs were no more robust to a test occluder condition than the DNNs trained without occlusion (i.e., poor generalization) while a score of 1 indicates that the DNNs were just as robust as those that received direct training on that test occluder condition. Using these measures, we found that natural occlusion-trained models attained normalized scores of .63 and .45 on artificial 1 and 2, respectively, while DNNs trained on artificial occluders resulted in normalized scores below 0.1 for natural occluders (**Figure 2b**). This pattern of results was highly consistent at a finer grain, emerging for each of the five DNN architectures that were investigated (**Figure 2c**). These findings suggest that applying more naturalistic occlusion during training leads to more generalized robustness in DNNs, and that artificial occluders used in certain computer vision applications (e.g., ^35,36^) might have little utility for real-world deployment where robustness to natural occlusion is critical.

To explore the relative contributions of natural occluder shape and natural occluder texture to these training benefits, we compared the performance of DNNs trained with artificial versus natural occluders. Importantly, the natural silhouette occluders were based on the same shapes as the natural occluders but shared the texture properties of the artificial occluders, thereby allowing us to disentangle these factors. Here, we made two observations in the normalized accuracies (**Figure 2b**). First, DNNs trained with artificial occluders showed poorer generalization to natural silhouette occluders in their normalized accuracies (0.57 and 0.74) than was observed for DNNs trained on natural silhouette when generalizing to artificial occluders (0.90 and 0.93). These findings suggest that training DNNs with more naturalistic occluder shapes promotes more generalized robustness to occlusion. Second, DNNs trained on natural silhouettes performed somewhat better when tested with natural occluders (0.16) in comparison to DNNs trained on artificial occluders (0.08 and 0.09), but these generalization benefits were asymmetric, as DNNs trained with natural occluders could generalize well to natural silhouettes (0.58). These findings indicate that a key advantage of training DNNs with natural occlusion arises from the textural properties of natural occluders. These patterns of results were evident for each of the DNN architectures (**Figure 2c**). Taken together, our results indicate that both the shape and textural properties of natural occluders promote generalized robustness in DNN models across a wide variety of occlusion conditions.

### Human alignment

To evaluate DNN-to-human alignment, we conducted a separate behavioral experiment that presented human observers with object images drawn from 8 different categories, which were applied with various types and intensities of occlusion (see **Figure 3**). We could then compare human classification accuracies across the various occluder types with the performance of DNNs models trained with different forms of occlusion, as shown in **Figure 4a**. A general finding was that overall classification accuracy was higher for the DNNs trained with either artificial occluders or natural silhouette occluders, as compared to the DNNs trained with natural occluders, presumably because this behavioral study evaluated recognition of objects presented behind uniformly white or black occluders. Of greater interest was the extent to which DNN models could predict human robustness to various forms of occlusion. This was evaluated by calculating the mean accuracy of each DNN model and human observer for a given occluder type and color (black or white), averaging across all occlusion intensities (20-90%), and then evaluated the relationship between these 18 accuracy scores across DNNs and humans (see **Methods**).

**Figure 3.**
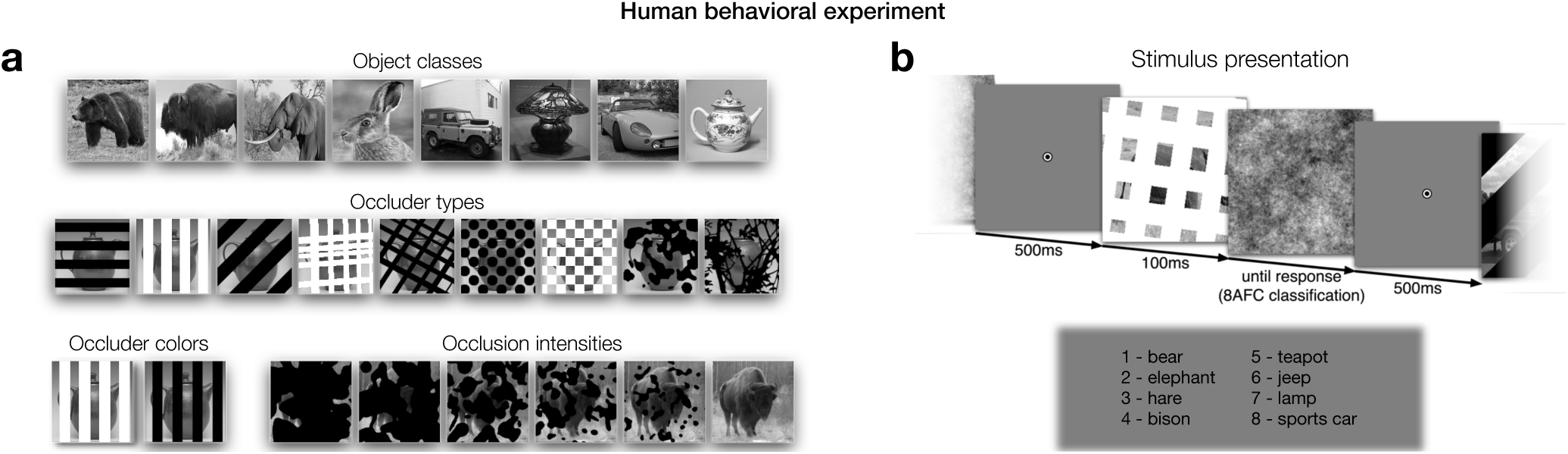
Design of human behavioral experiment, in which participants performed 8-AFC classification with occluded and non-occluded object images. (**a**) Example stimuli from the different object and occluder conditions. (**b**) Schematic of the stimulus presentation sequence. Images were rendered in grayscale to prevent participants from relying on basic color cues to perform the task. Short presentation times followed by backward masking with a Fourier phase-scrambled noise pattern was implemented to reduce the opportunity for higher-level cognitive processing.

**Figure 4.**
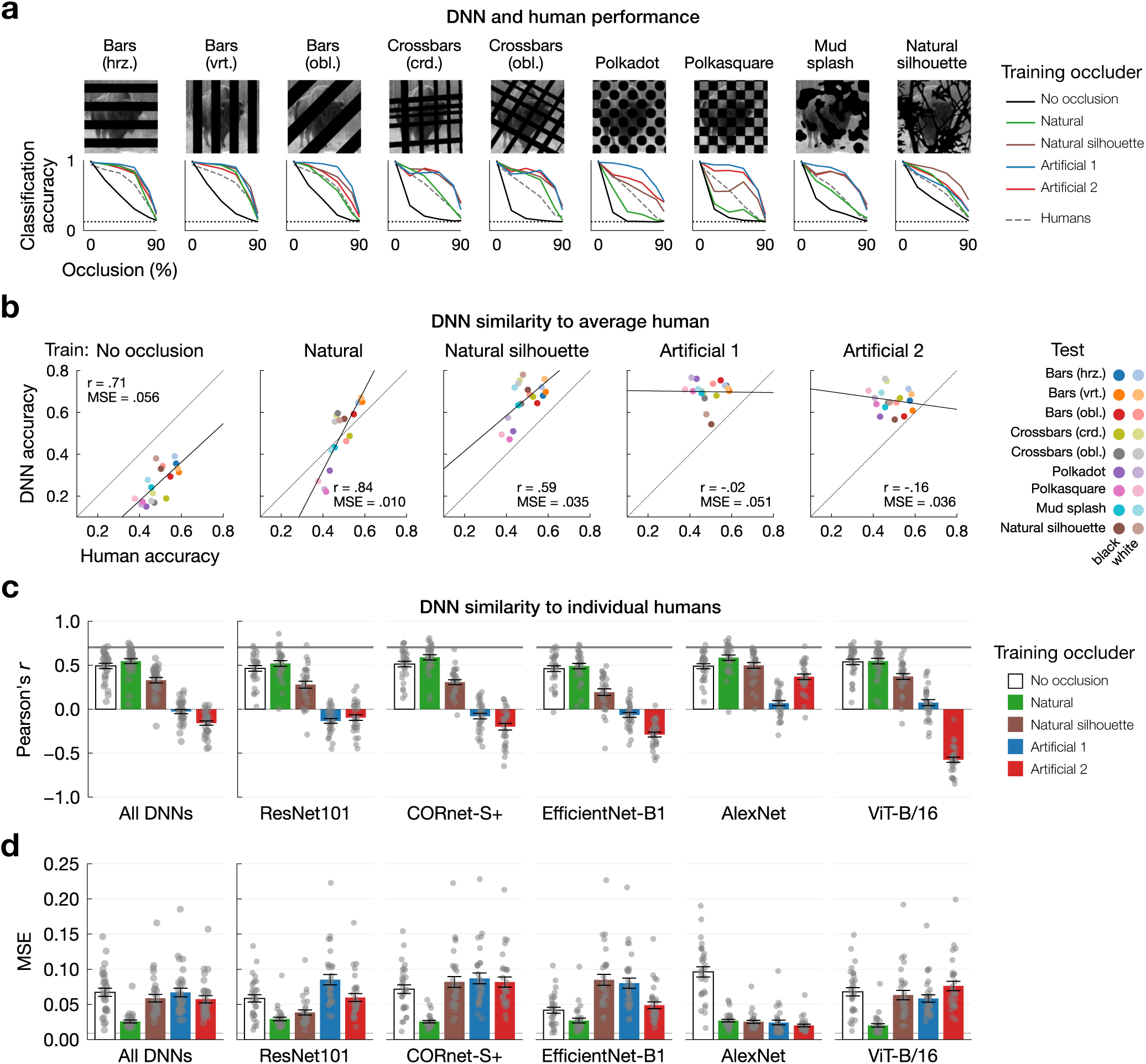
Human versus DNN performance during occluded object classification (dataset illustrated in Figure 3). (**a**) DNN and human performance (8-AFC classification accuracy) as a function of occlusion intensity, plotted separately for each occluder type. Scores are averaged across the eight object classes, two occluder colors, and human participants / DNN architectures. Horizontal dotted line indicates chance performance (0.125). (**b**) Scatterplots showing similarity between humans and DNNs trained with different occluder sets or without occlusion, each shown in a separate plot. DNN-human similarity was based on comparing the set of classification accuracies across 18 test conditions (nine occluder types and two occluder colors), indicated by the color of each point. Performance was averaged across the five DNN architectures and the 30 human participants prior to calculating the Pearson correlation and MSE between humans and DNNs. The solid line shows a linear model of the data and the dashed line indicates equal performance between DNNs and humans. (**c, d**) Bar plots showing DNN-human similarity when the same analyses were performed separately for individual human participants (gray points). Performance was calculated while aggregating scores across the five DNN architectures (left plot) and separately for each architecture (right plots). Bar height and error bars show the group mean and standard error, respectively, across human participants. The upper and lower bounds of the noise ceiling are spanned by the grey horizontal bar, calculated as the mean correlation between each participant and the remaining group (lower bound) and each participant versus the entire group (upper bound).

First, we compared the mean accuracies of humans with each set of DNN models trained on a particular occlusion augmentation (**Figure 4b**). For the occlusion-trained models, the Pearson correlation between humans and DNNs was highest for models trained with natural occluders (r = 0.84), followed by natural silhouette (r = 0.59), whereas DNNs trained with artificial occluders showed no positive relationship with human performance (r = −0.02 or −0.16). We performed this analysis at a finer grain by correlating individual human accuracy scores with the mean accuracies of each set of trained DNNs (**Figure 4c, left**) and further compared human scores with the accuracies of individual DNN models (**Figure 4c, right**). At each level of granularity, we observed a similar pattern of DNN-human similarity across the different training conditions. Specifically, natural occlusion training resulted in the highest similarity to humans, followed by training without occlusion, then natural silhouette, and finally the two artificial occluder conditions. While DNNs trained without occlusion were predictive of the relative differences in human accuracies across occluder types, these models were far less robust to occlusion than human observers, which was evident in our analyses of the mean-squared error (MSE) between DNN versus human accuracies (**Figure 4d**). Thus, taking into consideration both Pearson correlation and MSE measures of DNN-human similarity, it is evident that DNNs trained without occlusion are inadequate models of human performance.

We evaluated whether the observed differences in DNN-human correlation strength were statistically reliable by applying a Fisher transformation to the participant-wise correlation coefficients ^37^, which were then averaged across the models in each training condition and entered into a one-way, repeated-measures ANOVA. These analyses revealed a highly significant effect of training occluder set on DNN-human similarity (F(4,116) = 151.98, *p* < .0001). We then performed post-hoc, two-sided, pairwise t-tests to compare the different DNN training conditions, using Bonferroni-Holm correction for multiple comparisons ^38^. In comparison to the DNNs trained without occlusion, correlations with human performance accuracy were significantly higher for the DNNs trained with natural occluders (t(29) = 2.07, *p_corrected_* = .0466) and significantly lower for DNNs trained with artificial occluders (artificial 1: t(29) = 13.40, *p_corrected_* < .0001; artificial 2: t(29) = 19.29, *p_corrected_* < .0001), suggesting that exposure to natural and artificial occlusion had opposite effects on DNN-human alignment. Also, DNNs trained with natural silhouette occluders exhibited higher correlations with human performance than DNNs trained with artificial occluders (artificial 1: t(29) = 8.09, *p_corrected_* < .0001; artificial 2: t(29) = 17.61, *p_corrected_* < .0001), but lower correlations than DNNs trained with natural occlusion (t(29) = 12.73, *p_corrected_* < .0001) or without occlusion (t(29) = 6.03, *p_corrected_* < .0001), suggesting that, while both the texture and shape properties of natural occluders promoted DNN-human alignment, the benefits of natural shape alone were outweighed by the counteracting effect of artificial texture. Taken together, these findings suggest that by exposing DNNs to more naturalistic forms of occlusion, encompassing both shape and textural properties, one can attain improved DNN-human alignment in the processing of occluded objects.

### Control analyses

Given that the DNNs trained with artificial occluders were much more accurate overall in their classification of occluded objects (**Figure 4a,b**), we wondered whether this might be the cause of their deviations from human accuracy profiles. To address this question, we performed a control analysis in which DNN test performance was equated across the different training regimes, then recalculated DNN-human similarity. Starting with DNN models that were pre-trained without occlusion, we performed end-to-end fine-tuning with a given occlusion training regime for up to 4 training epochs until each model reached the mean accuracy of human performance on the test dataset (± 1 SEM). We used grid-searches to calibrate the learning rate appropriately for each DNN architecture, which was consistent across the different training conditions. Based on our prior finding that DNNs trained without occlusion show strong correlations with human accuracy while lacking in overall robustness, we anticipated that DNNs fine-tuned with fewer epochs of training with artificial occluders would show reduced levels of divergence from humans, but that differences between natural and artificial occlusion training would still persist.

After each DNN model was effectively calibrated to match average human accuracy (**Figure 5a**), we re-evaluated DNN-human similarity. **Figure 5b** shows the correlation between averaged DNN models and individual human accuracies for both the original and performance-equated DNNs. For performance-equated DNNs, condition-wise accuracies are plotted against those of the human participants (group mean) in **Figure 5c**.

**Figure 5.**
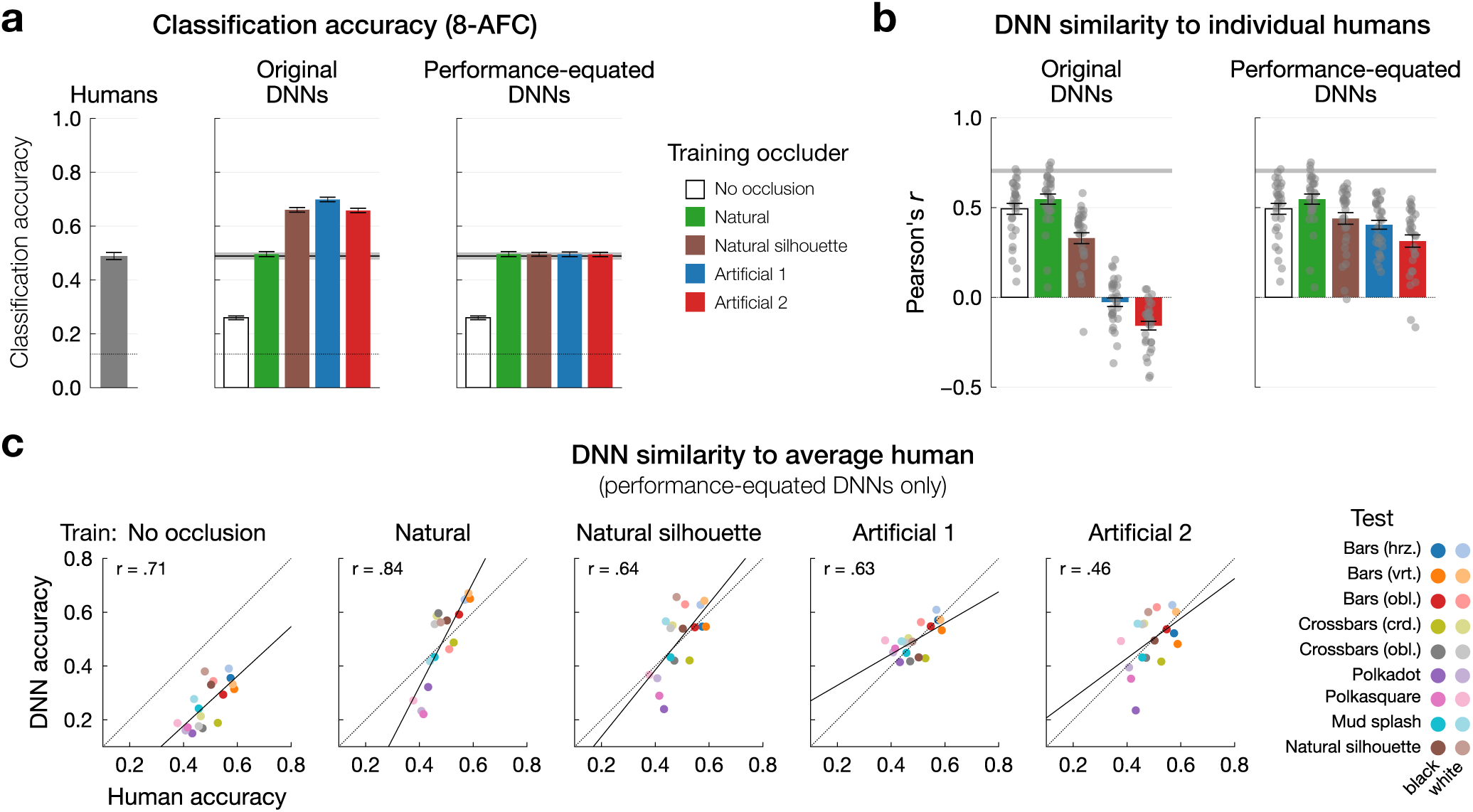
Control analyses exploring how DNN-human similarity might be explained by overall performance on the behavioral test dataset (see Figure 3). (**a**) Barplots showing 8-AFC classification accuracy on the behavioral test dataset across all images except for those without occlusion. The left plot shows human performance, with error bars showing SEM across participants (N=30). The middle plot shows performance for the original set of DNNs, averaged across the five model architectures, with error bars showing SD across the 30 subsets of images each shown to a different human participant. The right plot shows DNN performance after substituting a subset of the original DNNs with instances finetuned to attain human-level performance. For comparison, human performance is shown by the horizontal black (mean) and gray (SEM) bars. In all plots, the dotted line indicates chance performance (0.125) (**b**) Barplots showing DNN-human similarity (Pearson correlation of performance across conditions) for the original and performance-equated DNNs, calculated for each individual participant (gray points) and presented as in Figure 4c. (**c**) Scatterplots showing condition-wise accuracies for performance-equated DNNs versus humans (group mean). For corresponding scatterplots for the original set of DNNs, see Figure 4b.

Following the original analysis, the participant-wise correlation coefficients for the averaged models were Fisher-transformed ^37^, then subjected to a one-way, repeated-measures ANOVA, which revealed a strong, significant effect of occluder training condition (F(4,116) = 31.90, *p* < .0001). Post-hoc, two-sided, pairwise t-tests were then performed, again using Bonferroni-Holm correction for multiple comparisons ^38^. This analysis revealed that after DNNs were fine-tuned with artificial occluders, they exhibited significantly lower correlations with human performance than the original DNN models that were pre-trained without occlusion (versus artificial 1: t(29) = 4.24, *p_corrected_* = .0008; versus artificial 2: t(29) = 8.57, *p_corrected_* < .0001). Thus, DNN fine-tuning with artificial occluders leads to improvements in overall classification accuracy, but also introduces increasing divergence with behavioral patterns of human observers.

We also investigated how the DNN models performed after undergoing training with varying levels of occlusion severity. To do this, we evaluated instances of ResNet101 and EfficientNet-B1 that were trained on either weak (0-20%), moderate (0-50%) or strong (0-90%) levels of occlusion, and then tested these models on the behavioral test dataset. DNN accuracy and DNN-human similarity can be seen in **Figure 6**. The results revealed that natural occlusion training produced the most human-like profile of performance at each occlusion-training intensity, both in terms of correlation strength and mean squared error, in agreement with our previous results. We also observed that while stronger levels of occlusion training generally led to higher accuracy, there was no corresponding trend of further improvement in our correlational measure of DNN-human similarity. These results suggest that the emergence of human-like occlusion robustness in DNNs is much more tightly linked to qualitative aspects of occlusion exposure (i.e., natural vs. artificial) rather than quantitative aspects such as occlusion severity.

**Figure 6.**
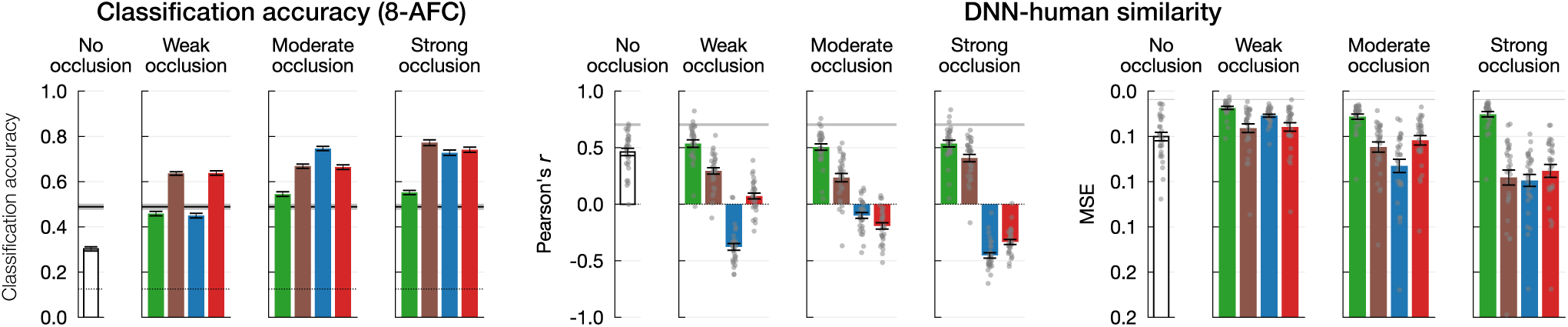
Performance of DNNs trained with different intensities of occlusion (ResNet101 and EfficientNet-B1 only). Occlusion intensities were 0-20%, 0-50% and 0-90% for weak, moderate and strong occlusion training, respectively. Left plot shows 8-AFC classification accuracy, presented as in Figure 5a. Middle and right plots show DNN-human similarity through both Pearson correlation and MSE across the condition-wise performance, presented as in Figure 4c,d.

### Robustness afforded by prior occlusion-training techniques

Next, we evaluated the robustness afforded by other occlusion-like image augmentations that are commonly used in computer vision, all of which relied on more artificial methods. We trained new sets of DNN models on each of three types of image corruption motivated by previous work. These included two configurations of RandomErase ^36^, in which training images are partially covered by rectangular occluders that are either filled with zero values or pixelated noise, and CutMix ^39^, in which each image in a training batch is occluded by a rectangular crop of another image in the batch. For CutMix, ground-truth labels were adjusted to reflect the relative size of the two different objects contained in each training image. We used the built-in PyTorch implementation of these dataset augmentations with default hyperparameters (https://docs.pytorch.org/vision/main/transforms.html). We found that DNNs trained using either RandomErase or CutMix performed poorly at generalizing to object images with artificial occluders, in comparison to our occlusion-trained DNN models (**Figure 7a, left**), and were likewise unable to generalize to objects with natural occluders. It is also worth highlighting that CutMix training involved the presentation of occluders with natural color and texture while lacking in the shape variations of our natural occluders. These findings point to the inadequacy of training DNNs with simple rectangular image crops as occluders to instill generalizable robustness to occlusion.

**Figure 7.**
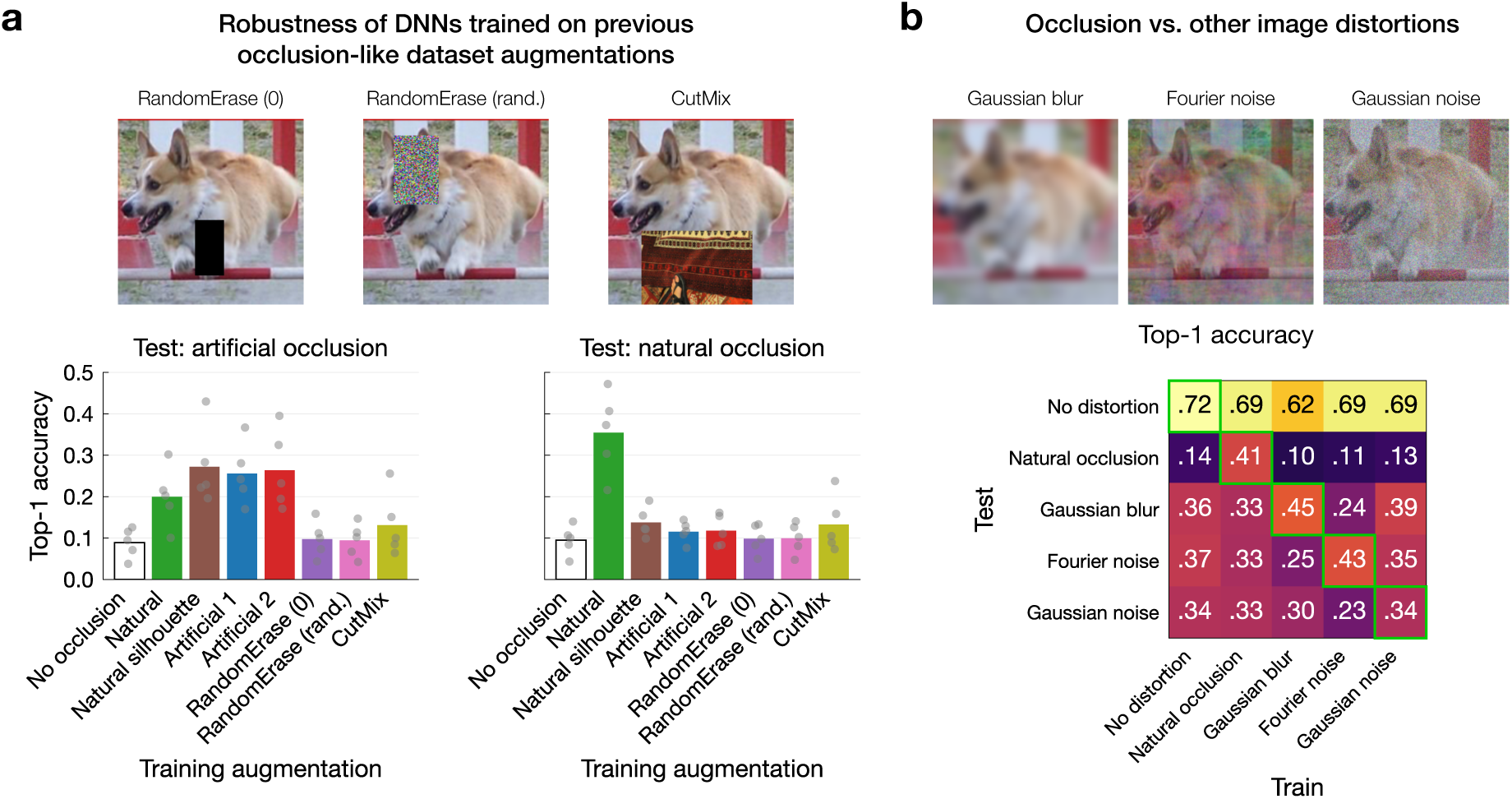
DNN robustness resulting from the present occlusion-training approach and other training dataset augmentations from the literature. Examples of each distortion type are shown at the top of each panel. All plots show 1000-way classification accuracy on the ImageNet-1K validation set applied with various image distortions. (**a**) Robustness to artificial and natural occluders from the Visual Occluders Dataset in DNNs trained with our approach versus other occlusion-like augmentations. Gray points show performance for each DNN architecture, with bar height indicating the mean across architectures. (**b**) Performance of DNNs (ResNet101) trained and tested on natural occlusion and other, non-occlusion image distortions. Green boxes indicate where trained and tested image distortions were of the same type.

### Occlusion versus other image perturbations

Finally, we explored how DNN robustness to occlusion might involve the acquisition of visual mechanisms are distinct from those that support robustness to other forms of image perturbation. Specifically, we asked whether DNNs trained with noisy or blurry images might show improved robustness to occlusion by evaluating cross-generalization performance across multiple trained models. To address this question, we trained further instances of ResNet101 with either strong Gaussian blur ^21^, Fourier or Gaussian noise ^20^, natural occlusion, or no distortion. Other image preprocessing steps were identical to that used for all previous models (see **Methods**) except that we no longer applied the modest levels of random blur (σ = 0.1 to 2 pixels) that were previously used for DNN training. Once trained, we evaluated each DNN on the ImageNet-1K validation set with each type of image corruption. Robustness to Gaussian and Fourier noise was evaluated at 9 different signal to signal-plus-noise ratios (SSNRs) following the implementation described in Jang et al. ^20^, after which we calculated an averaged classification accuracy score. For Gaussian blur, we used the ImageNet-C benchmark ^40^, taking the average accuracy across the five blur intensity levels (σ = 1-6 pixels). The results are presented in **Figure 7b**. Compared to the DNN trained with no image distortion, blur- and noise-trained DNNs were no more robust to natural occlusion (second row of the matrix) and the occlusion-trained DNN was no more robust to blur or noise (first two columns, last three rows). These results are consistent with the notion that occlusion presents a unique problem in real-world vision with its own characteristic solution space.

## Discussion

In the present study, we found that training DNNs with artificial versus naturalistic occlusion led to very different outcomes. DNNs trained with artificial occlusion greatly improved in their ability to recognize objects with artificial occluders, but remained vulnerable to naturalistic occlusion and showed much weaker correspondence with human vision. In comparison, models trained with natural occluders consistently showed a much more generalized, human-like form of robustness. We further found that the advantages of training with natural occlusion were partially reduced when the occluders were rendered as silhouettes during training, consistent with the notion that naturalistic aspects of occluder shape and texture are both important for promoting generalized robustness and DNN-human alignment. This pattern of results was strikingly consistent across different model architectures and manipulations of occlusion training intensity. Taken together, our findings provide novel evidence to support the ecological vision hypothesis ^41^, which assumes that human robustness to occlusion is acquired via learning from prevalent encounters with occluding foreground objects in natural vision. Thus, DNN models that are trained with naturalistic occlusion conditions will exhibit more human-aligned patterns of performance, even when tested with artificial occluders that deviate considerably from those viewed during training.

We found that natural and artificial occlusion training had opposite effects on human-alignment when referenced against standard training datasets. Specifically, natural occlusion training resulted in a more human-like performance profile whereas artificial occlusion training led to DNN model divergence. These findings run counter to the notion that human robustness to occlusion arises simply through frequent exposure to partial views of objects in a manner that can be dissociated from the specific visual properties of the occluder. Instead, our results suggest that humans acquire robustness to occlusion by learning to disentangle real objects that frequently obscure one another during natural vision. Previous theories have proposed that humans perceive partially occluded objects by relying on generic visual cues, such as ‘T’ junctions, which can be used to infer border-ownership, relative depth, and likelihood that two surfaces or object parts are likely to connect behind a foreground occluding object ^42–44^. Given that these cues are present under both natural and artificial occlusion, our results suggest that in addition to these mid-level visual properties for inferring how an implied object may continue behind an occluder, the visual system’s representation of higher-level shape, texture and other object properties (e.g., ^45^) may be critical for our ability to recognize objects under conditions of real-world occlusion.

In addition to superior DNN-to-human alignment, we found that DNN training with natural occluders led to generalized robustness to novel occluders outside of the training distribution, including artificially shaped occluders and occluders that lacked textural cues. In contrast, artificial occlusion-training conferred minimal advantage over standard training datasets when tested with out-of-distribution natural occluders. These findings expand upon previous reports in which artificial occlusion-training showed limited generalization when the same occluder was presented at novel image locations to DNNs ^46^. Given that DNNs trained with artificial occluders are unable to generalize well to natural occlusion, our findings raise serious concerns regarding previous studies that have relied on artificial occlusion methods to benchmark DNN robustness to occlusion (Naseer et al., 2021; Yun et al., 2019), as such metrics may lead to overinflated estimates of DNN’s ability to generalize to real-world occlusion. Going forward, our results call for more naturalistic dataset augmentations to both estimate and instill occlusion robustness in computational models.

While the artificial occluders that we developed here had uniform texture, other studies have applied non-uniform texture, such as pixelated noise ^31,36^ or natural image texture ^39,46^ to occluders during training. These techniques were not necessarily developed with the goal of inducing occlusion robustness *per se* ^but, see^ ^46^, but rather, as a regularization strategy for improving the classification, localization, or detection of non-occluded objects. Nevertheless, some of the above studies did ultimately conclude that training with artificial occlusion led to improved robustness to occlusion, but without testing for generalization to more realistic conditions of occlusion. Here, we showed that two of these approaches, namely CutMix ^39^ and RandomErase with pixelated noise ^36^, conferred minimal gains in robustness to both our artificial and natural occluders. Our findings suggest, counterintuitively, that the utility of these occlusion-training (or masking) techniques might be limited to visual tasks involving non-occluded objects.

We further found that when DNNs are trained with natural silhouette occluders, these models exhibit better generalization performance and superior human-alignment, in comparison to DNNs trained on artificial occluders, though they still fall short of DNNs trained with fully natural occluders. Our findings indicate that both occluder shape and texture contribute to the robustness of DNNs to natural occlusion.

In conclusion, we show that generalized, human-like recognition of occluded objects emerges in DNNs only when trained with naturalistic forms of occlusion. In comparison, DNN training with artificial forms of occlusion led to the acquisition of misaligned representations that failed to yield generalized robustness to natural occluders. Our findings suggest that the perceptual mechanisms underlying human occlusion robustness likely arise from learning to disentangle 3D objects that frequently overlap during natural human vision, in agreement with our ecological vision hypothesis. Our study further reveals that popular DNN training methods that rely on artificial occlusion techniques may face serious difficulties if subsequently tasked to generalize to the challenges of real-world occlusion. Finally, we presented multiple lines of evidence to suggest that real-world occlusion constitutes a unique problem space for visual learning, which can be distinguished from the learning of DNN mechanisms tailored to address other image challenges such as blur or visual noise corruptions.

## METHODS

### Visual Occluders Dataset

We created a dataset of occluder masks (256×256 pixels) that varied in their spatial form, texture, and the proportion of the image that they obscured (see **Figure 1**). The full dataset contained 32 different occluder types, 30 of which were computer-generated using geometric shapes or Fourier noise patterns, sometimes in combination with other image processing techniques, and then rendered as silhouettes. Two different sets of these artificial occluders were formed for this experiment (‘artificial 1’ and ‘artificial 2’), each of which contained eight of the base occluder types in the dataset. We also created natural occluders by segmenting objects from real-world photographs. A basis set of 772 photographs contained both natural and human-made objects (e.g., tree branches, flowers, window blinds, gates, etc.) and were a mixture of photographs taken by the present authors and images scraped from the internet. Their backgrounds were removed using Adobe Photoshop (https://www.adobe.com/products/photoshop.html) and were rendered with both their original image texture (‘natural’) or as silhouettes (‘natural silhouette’). For each occluder type, we generated sets of masks at 9 occlusion intensities between 10 and 90%, indicating the approximate area (±5 %) of the underlying image obscured by the occluder. For artificial occluder types, occlusion severity was directly controlled in the generation process, with a total of 1000 masks generated for each occluder type and intensity level. For the natural occluders, each occluder in the basis set was randomly scaled and cropped until it obscured the desired proportion of the image, or until 100 random crops had been performed without satisfying the intensity criterion. This yielded between 194 and 692 unique occluder masks per intensity level.

### DNN architectures

For this study, we evaluated a variety of different DNN architectures, including a vision transformer (*ViT-B/16* ^47^) and four convolutional neural networks (CNNs), three of which were feedforward (*ResNet101* ^1^, *AlexNet* ^48^, *EfficientNet-B1* ^49^) and one of which had local recurrent connections (*CORnet-S* ^8^). These architectures were selected due to their popularity and/or task-performance on Brain-Score (https://www.brain-score.org/), a composite benchmark that quantifies model alignment with primate neural activity and behaviors ^11^. The AlexNet, ResNet101, EfficientNet-B1, and ViT-B/16 architectures were obtained from the TorchVision library (v0.21.0), and were not altered. We made three modifications to the original CORnet-S architecture (referred to in other sections as CORnet-S+). First, we increased the number of features in the convolutional layers of the V1 block from 64 to 128. Second, the adaptive average pooling layer in the decoder block was altered such that it outputted feature maps of size 3×3×256 for each input image (previously 1×1×256). Third, the decoder module was changed from a single fully-connected layer to two fully-connected layers with a ReLU operation in between the two layers. These changes were implemented based on our preliminary studies to substantially improve transfer-learning after self-supervised training (these DNNs are not reported).

### DNN training procedures

All models were trained using PyTorch v2.2.0 to perform 1000-way object classification using the ImageNet-1K dataset ^50^, with a cross-entropy loss function. Training hyperparameters (see **Table 1**) followed version 1 of the PyTorch training recipe for each architecture (https://github.com/pytorch/vision/tree/main/references/classification) except for CORnet-S+, which followed the original paper ^8^. These were consistently applied across all instances of each architecture, irrespective of the dataset augmentation. Aside from occlusion, a common set of image pre-processing steps were performed based on well-documented procedures ^51^. These augmentations included randomized cropping/resizing, horizontal flipping, color jitter, Gaussian blurring (σ = 0.1 to 2.0 pixels), and image intensity normalization. We also reduced the range of the random resized crop, which had a default range of 0.08-1.0 in areal size, to an adjusted range of 0.8 to 1.0. This was done to make the DNN training procedure more tractable, as our preliminary studies indicated that the combination of occlusion and extreme image cropping proved too severe to allow for the task to be learned. For occlusion-trained DNNs, occlusion was applied between the randomized cropping/resizing and horizontal flipping. First, an occluder mask was chosen at random from corresponding set of training occluders (intensity levels: 0-50%). The occluder was then randomly flipped horizontally and vertically, each with 50% probability, then filled with a randomly chosen uniform color (except for natural occluders, which retained their original image texture). Finally, the occluder image was randomly cropped to 224×224 pixels, then superimposed onto the object image. At most, 80% of training images were occluded during training, to ensure that the DNNs retained strong classification performance for non-occluded objects.

**Table 1.** DNN training hyperparameters.

|  | *AlexNet* | *CORnet-S+* | *ResNet101* | *EfficientNet-B1* | *ViT-B/16* |
| --- | --- | --- | --- | --- | --- |
| epochs | 90 | 43 | 90 | 90 | 300 |
| batch size | 32 | 256 | 32 | 32 | 512 |
| optimizer | SGD | SGD | SGD | SGD | AdamW |
| learning rate | 0.01 | 0.1 | 0.1 | 0.1 | 0.003 |
| weight decay | 0.0001 | 0.0001 | 0.0001 | 0.0001 | 0.3 |
| momentum | 0.9 | 0.9 | 0.9 | 0.9 | - |
| warm-up LR scheduler | - | - | - | - | LinearLR<br>start_factor: 0.033<br>total_iters: 30 |
| main LR scheduler | StepLR<br>gamma: 0.1<br>step_size: 30 | StepLR<br>gamma: 0.1<br>step_size: 20 | StepLR<br>gamma: 0.1<br>step_size: 30 | StepLR<br>gamma: 0.1<br>step_size: 30 | CosineAnnealingLR<br>T_max: 270 |

Hyperparameters for each architecture were consistent across different dataset augmentations and learning objectives. Optimizers and schedulers indicate the names of the PyTorch v2.2.0 function used, with non-default settings shown below. SGD = stochastic gradient descent.

### Evaluating DNN robustness to occlusion

To evaluate robustness to occlusion, we measured top-1 classification accuracy on instances of the ImageNet-1K validation set applied with the occluders in the Visual Occluders Dataset. A separate instance of the entire validation set was generated with each occluder type and intensity, resulting in 288 occluded validation sets. We also evaluated DNNs on the validation set without occlusion. To create each image from the raw object and occluder datasets, object images were first resized such that the smaller dimension was 224 pixels, then center-cropped to 224×224 pixels. Occluders were then applied to each object image as described for the training images.

### Human behavioral experiment

To measure the similarity between DNN and human perception of occluded objects, we first conducted a behavioral experiment in which human participants classified images of objects occluded with a subset of occluders from the Visual occluders Dataset. Using the same images, we then obtained class predictions from each DNN and measured the correspondence between human and DNN performance across conditions.

#### Participants

30 human participants (20 female, mean age: 19.4 years, SD age: 1.22 years, age range: 18 to 50 years) were included in this experiment. All participants had normal or corrected-to-normal vision and no reported history of neurological disorder. Each participant provided written, informed consent, and received course credit or monetary compensation of $12 (USD). The study was approved by the Vanderbilt University Institutional Review Board (IRB 040945).

#### Stimuli

784 unique object images were used, with 752 used in the main experiment and 32 used in a practice session. Images were selected from 4 animate classes (bear, bison, elephant, hare) and 4 inanimate classes (jeep, lamp, sports car, teapot) contained in ImageNet-1K image database ^50^. These classes corresponded to the following ImageNet synsets, respectively: n02132136, n02410509, n02504458, n02326432, n03594945, n04380533, n04285008, n04398044. The 98 images for each class contained all 50 images from the validation set, with the additional 48 taken from the training set. Images were converted to grayscale, then resized so that the shortest dimension was 256 pixels, then center-cropped to 256×256 pixels. For each participant, this set of object images was pseudo-randomly allocated such that 32 images (4 per class) were shown without occlusion, and 8 images (1 per class) were shown with each unique combination of 9 occluder types (the entire artificial 1 occluder set plus natural silhouette), 2 occluder colors (black, white), and 5 occlusion intensities (20, 40, 60, 80, 90%, see **Figure 3a**). For each occluded image, an occluder mask was randomly selected from the pool of up to 1000 masks with the specified type and intensity in the Visual Occluders Dataset, then superimposed on the object. This random pairing of objects and occluders resulted in each subject viewing a unique set of 752 images, with 22560 total ‘unique’ images across the 30 participants (the actual number of unique images might be slightly lower due to overlap in the non-occluded condition and chance occurrences of repeated object-occluder pairings).

#### Procedure

Each participant viewed their designated subset of 752 stimuli presented in randomized order while performing an 8-alternative forced-choice classification task (see **Figure 3b**). The entire experiment took about 30 minutes to complete. The object images (10 × 10 degrees) were presented on an LED computer monitor at a viewing distance of 57 cm, with participants’ head position stabilized by a chin rest. Each trial began with a 500-ms fixation period, followed by the stimulus which was centered around the fixation point. After 100 ms, the stimulus was replaced by a pink Fourier-noise pattern, which remained on screen until the participant made a classification response. This ‘backwards-masking’ approach was used to minimize the opportunity for high-level top-down processing of the image, to thereby better capture ‘core’ object recognition performance ^52^. Participants indicated the object class by pressing one of 8 possible keys on the computer keyboard (4 keys per hand), with the mapping from keys to classes shown continuously below the stimulus window. To familiarize the participants with the task and the object-response key mapping, a short practice session was performed prior to the experiment, in which the 32 practice object images were shown without occlusion. The mapping of keys to object classes remained consistent across trials for a given participant, but was randomized across participants, with the constraint that all 4 animate objects were assigned to the fingers of one hand while the inanimate objects were assigned to the other hand.

### Evaluating DNN-human alignment

For each DNN, we first obtained 8-AFC class predictions for all 22560 unique stimuli from the behavioral experiment. To do this, images were resized from 256×256 pixels to 224×224 pixels and normalized to match the mean and standard deviation of the ImageNet-1K training image set (averaged across color channels) before being provided to the DNNs. We extracted the eight output values from the final layer corresponding to the test object classes, and then identified highest value to obtain an 8-AFC class prediction. We then calculated the mean classification accuracy for each of the 18 combinations of occluder type and color in the test dataset and compared this to the corresponding accuracies of each human participant using both Pearson correlation and mean squared error. Because each human participant observed a unique set of images, these condition-level similarity metrics allowed us to estimate human-to-human similarity and, in turn, contextualize DNN-to-human similarity against this theoretical maximum, i.e. the ‘noise-ceiling’. To calculate the noise-ceiling, we compared the performance of each participant to the mean performance across the remaining participants (lower bound) and across all participants (upper bound), following Representational Similarity Analysis from the neuroimaging literature ^53^. To ensure that DNN-to-human comparisons were mathematically consistent with the lower bound of the noise-ceiling, DNN performance was calculated based on images observed by the remaining group when being compared to each participant.

### Statistical analysis

To compare DNN-human similarity across different training conditions, Pearson correlation coefficients were calculated between the performance accuracy profiles of each DNN and each human participant of the behavioral experiment (N=30). Each performance profile was constituted by the 18 classification accuracies (8-AFC) for the 9 test occluder types and 2 test occluder colors). These coefficients were then Fisher-transformed ^37^ and averaged across the five different DNN architectures within each training condition. The resulting values were then entered into a one-way, repeated measures analysis of variance (ANOVA), which was implemented using the *statsmodels* Python package (v0.14.6, https://www.statsmodels.org). Post-hoc, two-tailed pairwise comparisons were performed using a Bonferroni-Holm correction across the 10 comparisons ^38^. Post-hoc tests were performed using the *pingouin* Python package (v0.5.5, https://pingouin-stats.org).

## Data availability

The source data and analysis code for reproducing all statistics and figures in this paper have been deposited in the following GitHub repository (github.com/ddcoggan/DNN_natural_occlusion). The codebase used to train DNNs is available at github.com/ddcoggan/model_trainer. The Visual Occluders Dataset and associated code can be obtained at github.com/ddcoggan/VisualOccludersDataset. All repositories are available under an MIT license.

## Acknowledgements

The authors would like to thank Kaylee Bashor for her technical assistance. This research was supported by the following grants from the National Institutes of Health, R01EY035157 and R01CA240274 to F.T. and P30EY008126 to the Vanderbilt Vision Research Center.

## Author contribututions

DDC and FT conceived and designed the study. DDC performed computational modeling, conducted the human behavioral experiment, and analyzed the data, with frequent guidance from FT. All manuscript drafts were written by DDC and edited by FT.

## Materials and correspondence

Correspondence should be addressed to Frank Tong or David Coggan. Material requests should be directed to David Coggan.

